# Optimized Multicolour Immunofluorescence Panel for Cattle T Cell Phenotyping by an 8-Colour, 10-Parameter Panel

**DOI:** 10.1101/2022.01.12.476027

**Authors:** Eduard O. Roos, William Mwangi, Wilhelm Gerner, Ryan Waters, John A. Hammond

**Affiliations:** The Pirbright Institute, Ash Road, Pirbright, Woking, Surrey, GU24 0NF, UK

**Keywords:** Flow cytometry, cattle PBMC, T cells, naïve T cells, effector memory T cells, central memory T cells, activated T cells, γδ T cells, T cell subsets

## Abstract

This multiplex staining panel was developed to differentiate cattle T cells into conventional (CD4 and CD8) and unconventional (γδ-TCR) subsets as well as their stage of differentiation and activation. The combination of CD45RO and CD62L allows the identification of naïve (T_Naïve_), central memory (T_CM_), effector memory (T_EM_) and terminal effector (T_TE_) T cells. Activated cattle T cells (T_AV_) can be identified by the cell surface expression of CD25. This panel was developed using cryopreserved cattle peripheral blood mononuclear cells (PBMCs) and tested on fresh as well as stimulated PBMCs. Therefore, this 8-colour, 10-parameter flow cytometry panel simultaneously identifies cattle T_Naïve_, T_AV_, T_CM_, T_EM_, T_TE_ and γδ-TCR cells. This panel will improve our ability to examine T cell response to pathogens and vaccines in cattle including the potential to identify previously undescribed subpopulations. Furthermore, this panel can be readily optimised for other bovid species as many of these reagents are likely to cross react.

## Background

Robust T cell responses are critical in the response to pathogen infection both for clearance and the formation of strong and broad memory responses (1). Cattle, like several other species, have a much higher proportion of γδ T cells compared to CD4 and CD8 (2–4). Consequently, it is important to study the entire T cell compartment simultaneously to fully characterise how immune protection arises and persists. Furthermore, as the research climate focusses on One Health approaches, the ability to study the immune response at high resolution in species that underpin global food security is essential.

Common to several non-model species, the first mAbs to study CD molecules on cattle T cells were derived from mouse immunizations with whole cattle PBMC populations or PBMC lysates. Antibodies were characterized in three international workshops on ruminant antigens (2,5,6). Together with identification of cross-reactive mAbs, this allowed the establishment of a basic toolbox to study cattle T cells and various subsets within them (7). However, several limitations still exist for the establishment of polychromatic flow cytometry staining panels. For example, many of the current 371 human CD molecules do not have an antibody that cross react with cattle. Another major limitation is the lack of useful mAbs that are labelled to a wider range of fluorochromes. This makes it difficult to expand panels beyond the three most common fluorochromes FITC (or AF488), PE and APC (or AF647). By conjugating existing T cell markers in-house we were able to develop a cattle T cell panel that utilises eight colours excluding PE and APC-conjugated antibodies. This allows the addition of specific antibodies, such as for cytokines or transcription factors, that maximises the broader utility of this panel for individual research needs. Additionally, if more of the available mAbs would be conjugated to fluorochromes that are excited by the violet laser, the panel can be further expanded.

This OMIP identifies all three main cattle T cell subsets (CD4, CD8 and γδ), as well as their subsets that are activated (T_AV_) or in the distinct differentiation states of naïve (T_Naïve_), central memory (T_CM_), effector memory (T_EM_) and terminal effector (T_TE_). The gating strategy we used initially identifies the two αβ T cell subsets CD4 (mAb clones CC8/CC30) and CD8 (mAb clone CC63) as well as the γδ T cells (mAb clone GB21A) (2,8) (Fig. 1). Like in swine and chickens, γδ T cells constitute a major T cell subset in cattle blood and can comprise more than 50% of circulating T cells (3,9). To identify activated T cells, CD25 (mAb clone IL-A111) can be used (8,10–12), whereas the memory state of the cells can be defined using the CD45RO (mAb clone IL-A116) and CD62L (mAb clone CC32) cell surface markers (6,8, 13-15) (Fig. 1). Using this gating strategy, the following known subsets can be identified for the helper T cells, T_Naïve_ (CD3^+^γδ-TCR^-^CD4^+^CD25^-^CD45RO^-^CD62L^+^), T_CM_ (CD3^+^γδ-TCR^-^CD4^+^CD25^-^CD45RO^+^CD62L^+^), T_EM_ (CD3^+^γδ-TCR^-^CD4^+^CD25^-^CD45RO^+^CD62L^-^), T_TE_ (CD3^+^γδ-TCR^-^CD4^+^CD25^-^CD45RO^-^CD62L^-^) and T_AV_ (CD3^+^γδ-TCR^-^CD4^+^CD25^+^). Similarly, the cytotoxic T cells can be separated into T_Naïve_ (CD3^+^γδ-TCR^-^CD8α^+^CD25^-^CD45RO^-^CD62L^+^), T_CM_ (CD3^+^γδ-TCR^-^CD8α^+^CD25^-^CD45RO^+^CD62L^+^), T_EM_ (CD3^+^γδ-TCR^-^CD8α^+^CD25^-^CD45RO^+^CD62L^-^), T_TE_ (CD3^+^γδ-TCR^-^CD8α^+^CD25^-^CD45RO^-^CD62L^-^) and T_AV_ (CD3^+^γδ-TCR^-^CD8α^+^CD25^+^). Furthermore, γδ T cells can also be identified by (CD3^+^γδ-TCR^+^) (Fig. 1; Online Table 3). A major improvement by this panel is the simultaneous analysis of all these T cell subsets in a single sample, hence reducing the variation between replicates and the number of samples needed per animal.

**Figure 1.**
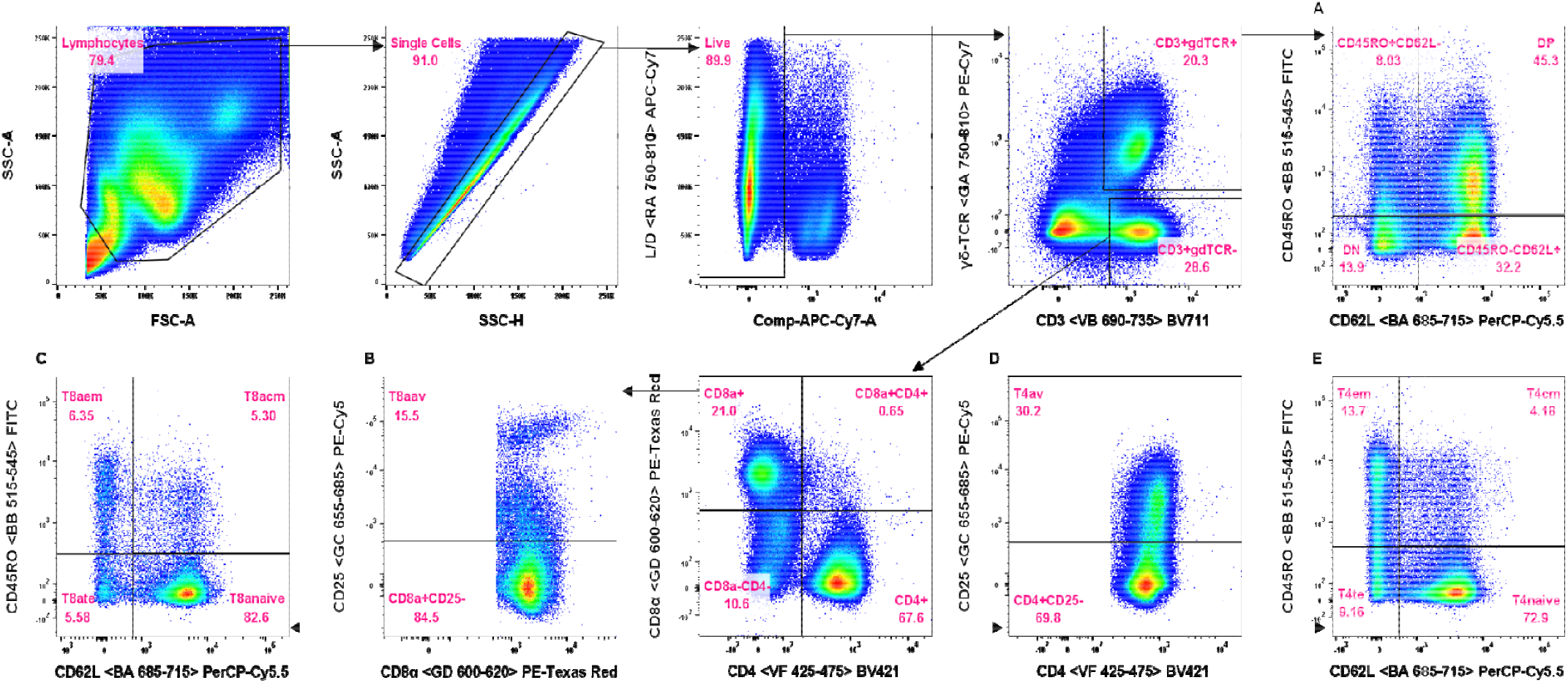
Gating strategy of the cattle T-cell panel into CD4^+^, CD8^+^ and γδ-TCR^+^ subsets. A) Sub-gating of the CD3^+^γδ-TCR^+^ subset with CD45RO and CD62L. B) Sub-gating of the CD3^+^ γδ-TCR^-^CD8α^+^subset with CD25. C) Sub-gating of CD3^+^ γδ-TCR^-^CD8α^+^CD25^-^ subset with CD45RO and CD62L. D) Sub-gating of CD3^+^ γδ-TCR^-^CD4^+^ with CD25. E) Sub-gating of CD3^+^ γδ-TCR^-^CD4^+-^CD25^-^ subset with CD45RO and CD62L. Sub-gating of the CD4 and CD8α with CD45RO and CD62L allows the identification of effector memory (T_EM_), central memory (T_CM_), terminal effector (T_TE_) and naïve (T_Naïve_) T cells. The CD25 identifies the activated T (T_AV_) cells, that include the T regulatory cells.

The panel was designed and optimised on a BD LSRFortessa and was tested on a BD Aria IIIU. The BD Aria IIIU allows for sorting of the T cell subsets. Further adaptations to the panel are enabled by having both the PE and APC channel empty, for which many antibodies are commercially available. If more reagents become available in the violet channel, they can easily be added to the panel with only minor influence on compensation requirements.

In conclusion, this cattle T cell panel will advance the understanding of the cattle immune response as it allows the measurement of all major T cell subsets and their differentiation stage within a single sample.

## Similarity to published OMIPs

None to date.

## Acknowledgements

The authors wish to acknowledge the valuable input of Dr Katy Moffat for technical support on the instruments. This project received funding from the European Union’s Horizon 2020 research and innovation programme under the VetBioNet grant agreement (731014). The authors would like to acknowledge the Pirbright Flow Cytometry facility and the Immunological Toolbox unit supported by the United Kingdom Research and Innovation-Biotechnology and Biological Sciences Research Council awards (BBS/E/I/00007038 and BBS/E/I/00007039); JAH and WG were further supported by BBS/E/I/00007030 and BBS/E/I/00007031.

