## Supplementary material for "Optimized Multicolour Immunofluorescence Panel for Cattle T Cell Phenotyping by an 8-Colour, 10-Parameter Panel": Online supporting information

**Online supporting information Optimized Multicolour Immunofluorescence Panel:** Cattle T Cell Phenotyping by an 8-Colour, 10-Parameter Panel

**Online Table 1. Instrument configuration.**

| Laser  Wavelength (nm) | Laser Power (mW) | Laser Type | Detector | Spectral Range for Detector (nm) | Dichroic LP Filter (nm) | Band Pass (nm) | Fluorochromes used |
| --- | --- | --- | --- | --- | --- | --- | --- |
| 640 (Red – R) | 40 | COHERENT  CUBE 640-40C | RA750-810 | 750-810 | 750 | 780/60 | Live/Dead NIR |
|  |  |  | RB710-753 | 710-753 | 710 | 730/45 | None |
|  |  |  | RC663-677 | 663-677 | --- | 670/14 | None |
| 561 (Green – G) | 50 | COHERENT Sapphire 561 LP | GA750-810 | 750-810 | 750 | 780/60 | PE-Cy7 |
|  |  |  | GB685-735 | 685-735 | 685 | 710/50 | PE-Cy5.5 |
|  |  |  | GC655-685 | 655-685 | 635 | 670/30 | PE-Cy5 |
|  |  |  | GD600-620 | 600-620 | 600 | 610/20 | PE-Texas Red |
|  |  |  | GE575-590 | 575-590 | --- | 582/15 | None |
| 488 (Blue – B) | 50 | COHERENT Sapphire 488 LP | BA685-715 | 685-715 | 685 | 695/40 | PerCP-Cy5.5 |
|  |  |  | BB515-545 | 515-545 | 505 | 530/30 | FITC |
| 405 (Violet – V) | 50 | COHERENT  CUBE 405-50C | VA750-810 | 750-810 | 750 | 780/60 | None |
|  |  |  | VB690-735 | 690-735 | 690 | 710/50 | BV711 |
|  |  |  | VC650-670 | 650-670 | 635 | 660/20 | None |
|  |  |  | VD595-620 | 595-620 | 595 | 610/20 | None |
|  |  |  | VE500-550 | 500-550 | 475 | 525/50 | None |
|  |  |  | VF425-475 | 425-475 | --- | 450/50 | BV421/DyLight405 |

The Optimized Multicolor Immunofluorescence Panel was optimized on a LSRFortessa cytometer (BD Biosciences). The cytometer had a four-laser configuration, with the listed optical configuration. The long pass (LP) filter allows the longer wavelengths through to the detector and reflects the shorter wavelengths down the optical path.

| **Specificity** | **Clone** | **Fluorochrome** | **Vendor** | **Catalogue #** | **Dilution** | **Time** | **Step** |
| --- | --- | --- | --- | --- | --- | --- | --- |
| anti-Mouse IgG1 | A85-1 | BV711 | BD | 565786 | 1:3000 | 15’ | Second |
| Biotin | Streptavidin | BV421 | BioLegend | 405226 | 1:3000 | 15’ | Fourth |
| Dead cells | - | APC-Cy7 | Thermo Fisher Scientific | L10119 | 1:2000 | 10’ | Last |

**Online Table 2.** Commercial reagents used in OMIP-XXX

**Online Table 3.** In-house conjugated reagents used in OMIP-XXX

| **Antibody** | | | | **Labelling kit** | | | **Staining conditions** | | |
| --- | --- | --- | --- | --- | --- | --- | --- | --- | --- |
| **Specificity** | **Clone** | **Vendor** | **Catalogue #** | **Fluorochrome** | **Vendor** | **Catalogue #** | **Dilution used for staining** | **Time** | **Step** |
| CD8α | CC63 | ITBX^a^ | ITB00316 | PE-Texas Red | Bio-Rad | LNK172PETR | 1:4000 | 15’ | Third |
| CD45RO | IL-A116 | ITBX | ITB00286 | FITC | Bio-Rad | LNK061F | 1:2000 | 15’ | Third |
| CD62L | CC32 | ITBX | ITB00342 | PerCP-Cy5.5 | Bio-Rad | LNK142PERCPCY5.5 | 1:4000 | 15’ | Third |
| CD25 | IL-A111 | ITBX | ITB00283 | PE-Cy5 | Bio-Rad | LNK082C | 1:4000 | 15’ | Third |
| γδ-TCR | GB21A | Kingfisher Biotech | WSC0578B-100 | PE-Cy7 | Bio-Rad | LNK112PECY7 | 1:2000 | 15’ | Third |

^a^Immunological Toolbox (1)

**Online Table 4.** In-house biotinylated reagents used in OMIP-XXX

| **Antibody** | | | | **Labelling kit** | | **Staining conditions** | | |
| --- | --- | --- | --- | --- | --- | --- | --- | --- |
| **Specificity** | **Clone** | **Vendor** | **Catalogue #** | **Vendor** | **Catalogue #** | **Dilution used for staining** | **Time** | **Step** |
| CD4 | CC8 or CC30 | ITBX^a^ | ITB00321 or ITB00340 | Thermo Fisher Scientific | A39257 | 1:1000 | 15’ | Third |

^a^Immunological Toolbox (1)

**Online Table 5.** Primary antibodies used in OMIP-XXX

| **Antibody** | | | | | **Staining conditions** | | |
| --- | --- | --- | --- | --- | --- | --- | --- |
| **Specificity** | **Clone** | **Isotype** | **Vendor** | **Catalogue #** | **Dilution used for staining** | **Time** | **Step** |
| CD3 | MM1A | IgG1 | Bio-Rad | MCA6080 | 1:1000 | 15’ | First |

| Antigen | Clone | Isotype | Version 1^b^ | Version 2^c^ | Version 3^d^ |
| --- | --- | --- | --- | --- | --- |
| CD3 | MM1A | IgG1 | FITC | PE-Cy7 | BV711 |
| CD4 | CC8/CC30^a^ | IgG2a/1 | DyLight405 | DyLight405 | BV421 |
| CD8α | CC63 | IgG2a | PE-Tex Red | PE-Tex Red | PE-Tex Red |
| CD45RO | IL-A116 | IgG3 | AF700 | FITC | FITC |
| CD62L | CC32 | IgG1 | PerCP-Cy5.5 | PerCP-Cy5.5 | PerCP-Cy5.5 |
| CD25 | IL-A111 | IgG1 | PE-Cy5 | PE-Cy5 | PE-Cy5 |
| γδ-TCR | CC105/GB21A | IgG2b | PE-Cy7 | AF700 | PE-Cy7 |
| L/D NIR | N/A | N/A | APC-Cy7 | APC-Cy7 | APC-Cy7 |

**Online Table 6.** Different versions of the panel as it underwent changes in antibody and fluorochrome combinations.

Summary of changes made to optimise the panel:

^a^Use of clone CC30 should be the preferred option as discussed in Hints and tips section below.

^b^Version 1: Original panel design.

^c^Version 2: Moved CD3 from FITC to PE-Cy7; CD45RO moved to FITC; γδ-TCR moved to AF700. Changed the clone from CC105 (WC1) for the γδ-TCR to GB21A.

^d^Version 3: Moved CD3 from PE-Cy7 conjugated antibody to a primary secondary stain with BV711 as secondary; γδ-TCR moved from AF700 to PE-Cy7; CD4 changed to BV421 as a biotin-streptavidin staining step.

^a^Use of clone CC30 should be the preferred option as discussed in Hints and tips section below.

| **Subset** | **Phenotype** | **References** |
| --- | --- | --- |
| **Helper T Cells** |  |  |
| Naïve T cells | CD3^+^γδ-TCR^-^CD4^+^CD25^-^CD45RO^-^CD62L^+^ | (2–5) |
| Central Memory T cells | CD3^+^γδ-TCR^-^CD4^+^CD25^-^CD45RO^+^CD62L^+^ | (3,5,6) |
| Effector Memory T cells | CD3^+^γδ-TCR^-^CD4^+^CD25^-^CD45RO^+^CD62L^-^ | (3,5,6) |
| Terminal Effector T cells | CD3^+^γδ-TCR^-^CD4^+^CD25^-^CD45RO^-^CD62L^-^ | (7–10) |
| Activated T cells | CD3^+^γδ-TCR^-^CD4^+^CD25^+^ | (11,12) |
| **Cytotoxic T cells** |  |  |
| Naïve T cells | CD3^+^γδ-TCR^-^CD8α^+^CD25^-^CD45RO^-^CD62L^+^ | (2–5) |
| Central Memory T cells | CD3^+^γδ-TCR^-^CD8α^+^CD25^-^CD45RO^+^CD62L^+^ | (3,5,6) |
| Effector Memory T cells | CD3^+^γδ-TCR^-^CD8α^+^CD25^-^CD45RO^+^CD62L^-^ | (3,5,6) |
| Terminal Effector T cells | CD3^+^γδ-TCR^-^CD8α^+^CD25^-^CD45RO^-^CD62L^-^ | (7–10) |
| **γδ T cells** |  |  |
| γδ-TCR cells | CD3^+^γδ-TCR^+^ | (13) |

**Online Tale 7.** Cattle T cell subsets of major importance.

**Online Table 8.** Antibodies tested but not used for the final panel.

| **Specificity** | **Fluorochrome** | **Antibody clone** | **Vendor** | **Dilution** | **Reason for exclusion** |
| --- | --- | --- | --- | --- | --- |
| WC1 | PE-Cy7 | CC105 | ITBX^b^ | 1:1000 | GB21A clone detects all γδ-TCR whereas CC105 clone only detects the WC1^+^ subset of the γδ-TCR. Thus, CC105 was replaced with GB21A. No available fluorochromes to include as a conjugated antibody in this panel. |
| WC1.1 | AF488 | CC101 | ITBX | 1:10 | No available fluorochromes to include as a rapid conjugated antibody in this panel. |
| WC1.2 | AF488 | CC115/CC117 | ITBX | 1:10 | No available fluorochromes to include as a rapid conjugated antibody in this panel. |
| CD4 | BV421 | CC30 | ITBX | 1:1000 | Staining profile similar to CC8 in original experiments. Though CC30 should be the preferred option (explained in Hints and tips section below). |
| IFN-γ | PE | CC302 | ITBX | 1:2000 | Only used to demonstrate cytokine staining and flexibility of panel (explained in Hints and tips section below). |

^a^Alexa Fluor 488 was a Thirdary whole anti-IgG antibody used at 1:2000.

^b^ITBX – Immunological Toolbox (<https://www.immunologicaltoolbox.co.uk/>; visited: 04/10/2021).

**Developmental Strategy**

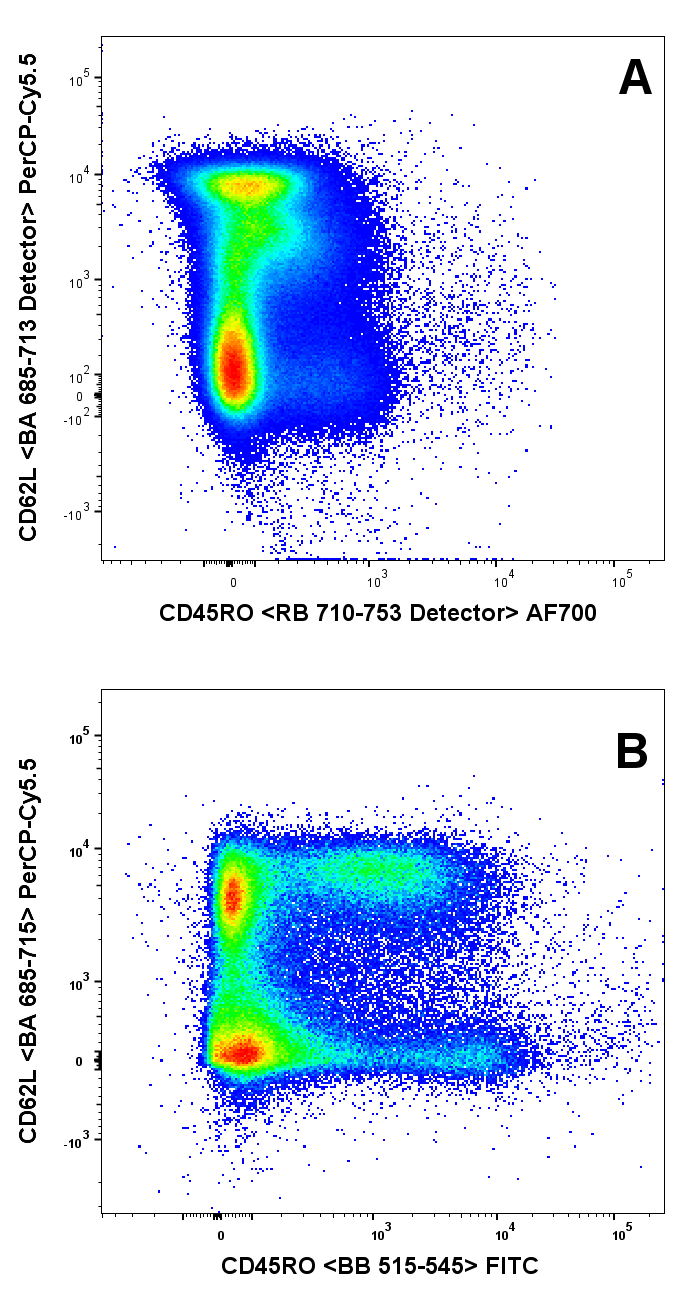

**Online Figure 1**

Separation of CD45RO on AF700 (A, Version 1) versus CD45RO on FITC (B, Version 3). The dot plots represent all live cells that were acquired for each experiment.

The 8-colour T cell panel was designed for a BD Fortessa LSR with the configuration as outlined in the Online Table 1. We selected mouse anti-cattle monoclonal antibodies (mAb) that are available from the [UK Immunological Toolbox](https://www.immunologicaltoolbox.co.uk/) (1). The panel identifies different T cell subsets with CD4 (helper) and CD8 (cytotoxic) T cells such as naïve (T_Naïve_), central memory (T_CM_), effector memory (T_EM_), terminal effector (T_TE_) and, activated T cell subsets as well as cells expressing the cattle γδ T cell receptor (γδ-TCR) (Fig. 1, Online Table 3).

The mAbs are not commercially available in the desired fluorochrome combinations, due to the limited availability of fluorochrome-conjugated mAbs for veterinary species. As most anti-cattle mAbs are only available as FITC (or AF488), PE or APC (or AF647) from Bio-Rad and Thermo Fisher Scientific and just as purified mAbs from most other vendors.

The mAbs were conjugated to the desired fluorochromes using commercially available conjugation kits (we refrained from custom conjugations to ensure accessibility to other laboratories) (Table 2). The in-house conjugation kits that were used are the LYNX range (Bio-Rad) and the EZ-Link Sulfo-NHS-LC-Biotin (Thermo Scientific) as listed in Table 2.

*Version 1*

The first version of the panel was not able to separate the CD45RO on AF700 (RA 710-753 detector) to a desirable level as was achieved by switching the CD45RO to FITC (Online Figure 1). This was the main concern for this version of the panel as all the other markers were able to discriminate their intended surface markers with the desired resolution. However, the mAb CC105 (Online Table 2), that recognises WC1 (workshop cluster 1) on the γδ-TCR cells was replaced by GB21A which sees all the γδ-TCR cells as opposed to the WC1 subset (13).

*Version 2*

Switching the CD45RO from AF700 (RA 710-753 detector) to FITC (BB 515-545 detector) meant that the CD3 had to be changed. The CD3 was moved to PE-Cy7 (GA 750-810 detector) to try and improve its resolution, which in turn necessitated moving the γδ-TCR to AF700 (RA 710-753 detector). Although the CD3 and γδ-TCR had an acceptable resolution both could still be improved (Online Figure 2).

*Version 3*

The unsatisfactory separation of the γδ-TCR on AF700 (RA 710-753 detector) required that we revert it back to PE-Cy7 (GA 750-810 detector) (Online Figure 2) and as a result the CD3 had to be moved to a different channel. This led to a different staining strategy as the CD3 had a very low resolution as a conjugated Antibody on the other channels (FITC and PE-Cy7 dilution were already at 1:62.5 and 1:125, respectively). A primary and secondary antibody staining step was considered, which allowed the use of secondaries that are available in the violet channels. The CD3 was subsequently moved to the BV711 (VB 750-810 detector) channel and had an optimal separation (Online Figure 2). The option to move the CD4 to BV421 (VF 425-475 detector) as a biotin-streptavidin stain greatly improved the mean fluorescence intensity of the mAb and resulted in a better resolution of the helper T cells (Online Figure 3). After these changes to the panel all the populations could be resolved satisfactorily.

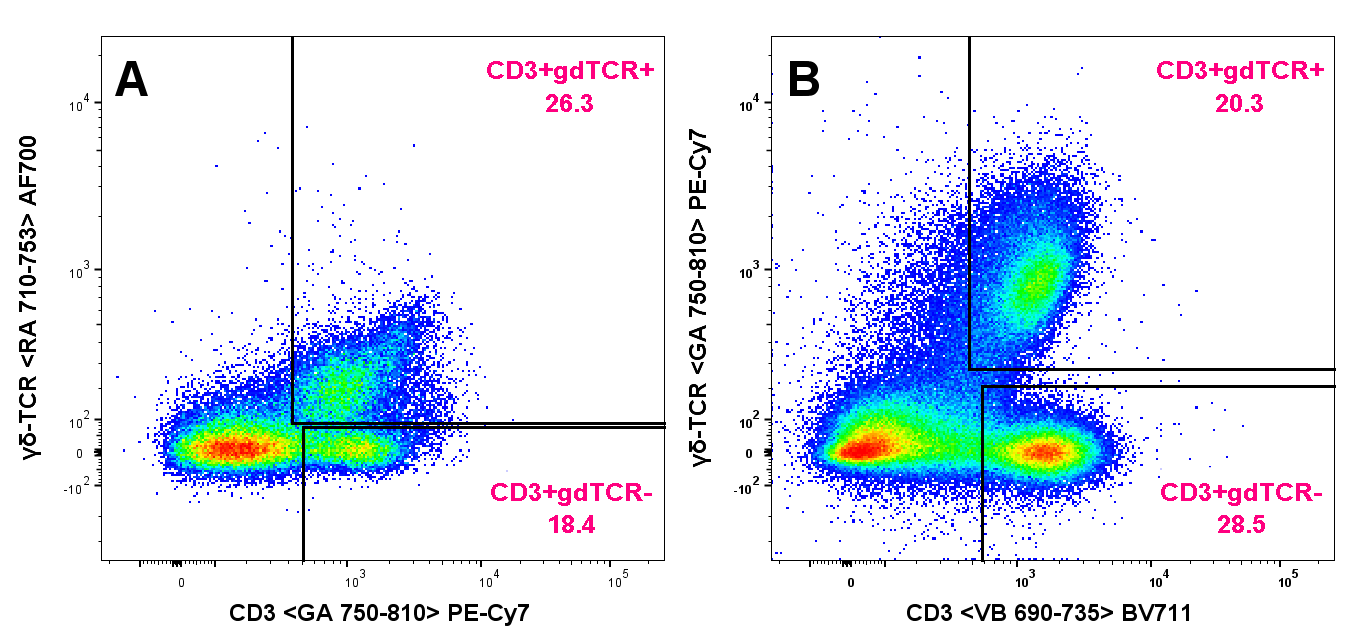

**Online Figure 2**

Separation of CD3 and γδ-TCR in Version 2 (A) versus Version 3 (B). The dot plots represent all live cells that were acquired for each experiment.

*Hints and tips*

**Online Figure 3**

Separation of CD4 on DyLight 405 (A) and BV421 (B). The dot plots represent all live CD3^+^ cells.

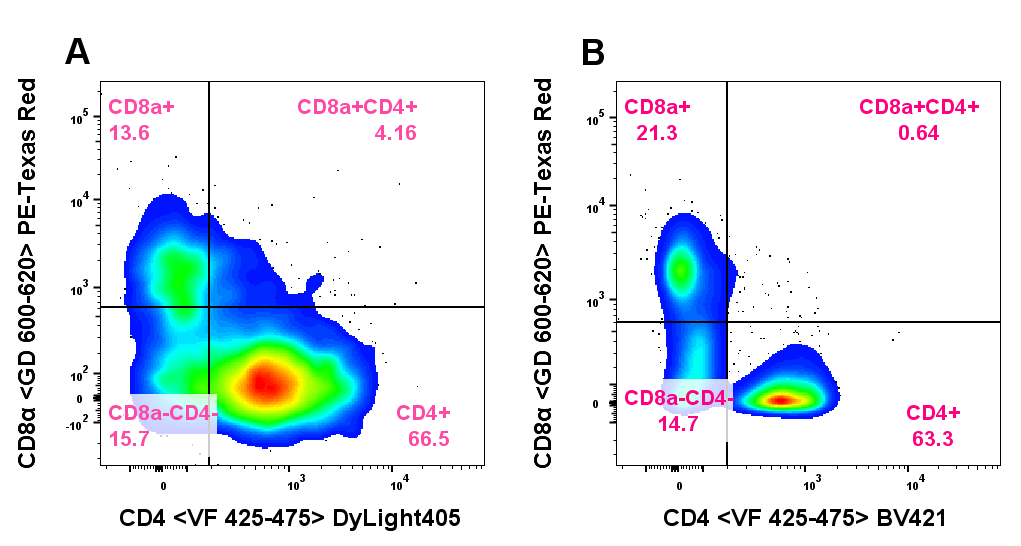

As outlined by previous research (14–16), the selection of the CD4 clone is a critical point. In this panel we initially used mAb clone CC8, but when this panel was tested in a larger number of animals, we came across a non-reactive animal for this clone. We tested mAb clone CC30 instead of CC8 and CD4 detection could be restored in this animal (Online Figure 4).

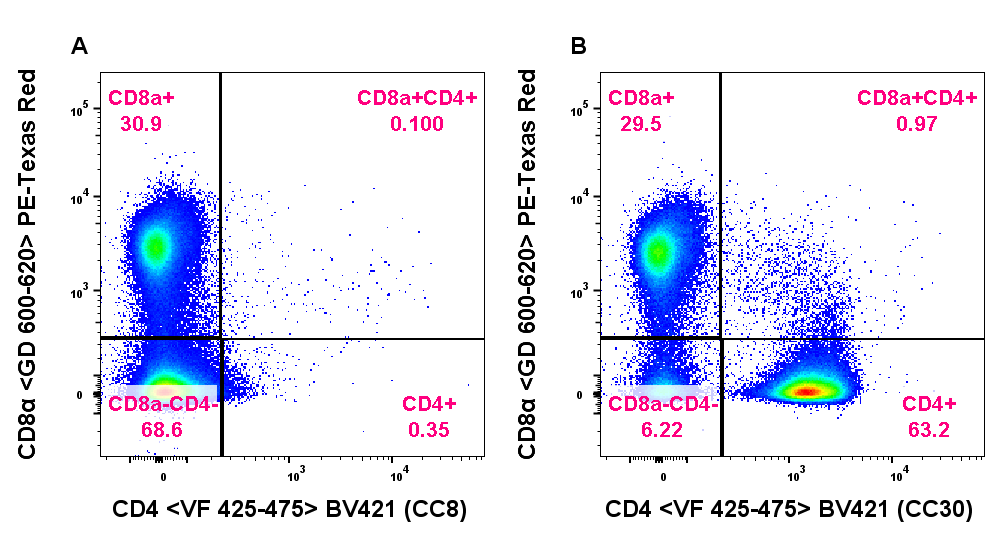

**Online Figure 4**

Comparison of two CD4 mAb clones, CC8 (A) and CC30 (B) in a cattle presumingly expressing a CD4 variant. Plots show CD3^+^γTCR^-^ pre-gated cells.

Both the PE and APC channels have purposefully been left unused to allow the incorporation of other antibodies that are often commercially available in only these two fluorochromes. This also allows the addition of PE or APC-conjugated mAbs specific for cytokines into the panel, since these bright fluorochromes are ideal for intracellular cytokine staining. An example of such a cytokine staining using this panel with anti-IFN-γ mAb clone CC302, conjugated with PE, is shown in Online Figure 5.

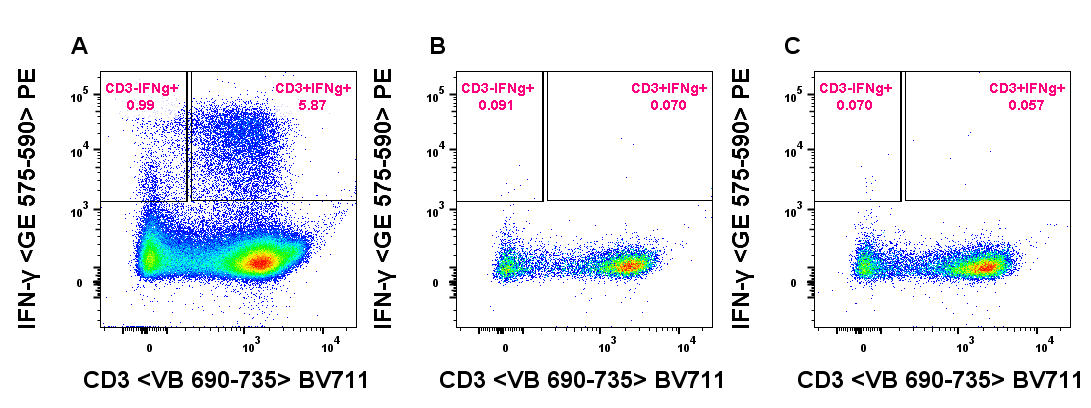

**Online Figure 5**

IFN-γ staining in addition to the 8-colour staining described in this OMIP. Cells were gated on live singlets (not shown). IFN-γ in PE (y-axis) versus CD3 (BV711) on the x-axis. Three different conditions are indicated: (A) PMA/Ionomycin + Brefeldin A (BFA) (B) BFA only (C) without PMA/Ionomycin and BFA.

Determining whether staining the Live/Dead at the end of our staining (done to include all dead cells) protocol had any influence on the read out the Full stain, unstained and single colour Live/Dead were compared (Online Figure 6). The results indicate that there is no difference if the Live/Dead NIR is included as the final staining step.

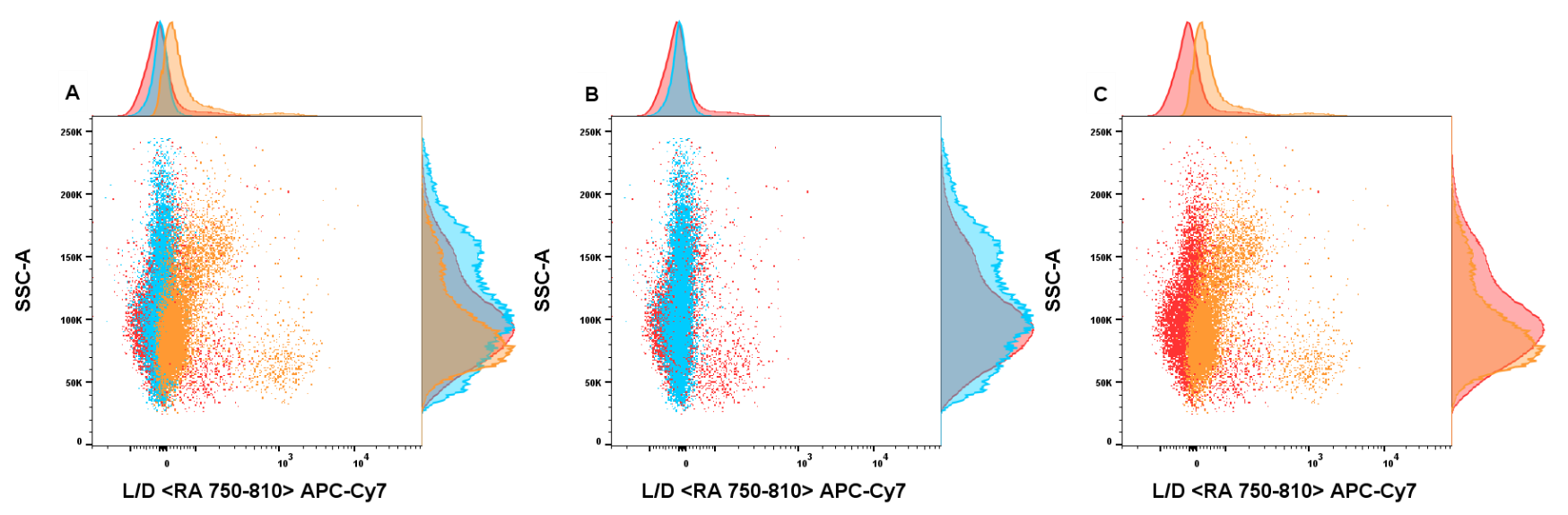

**Online Figure 6**

To determine the influence of the amine reactive Live/Dead NIR dye a single colour stain, fluorescence minus one (FMO) and full stain sample were used. A) All three conditions superimposed with the full stain (Red), FMO (blue) and single colour stain (Orange). B) Comparison of the Live/Dead NIR FMO overlayed on the full stain. C) Single colour stain overlayed on the full stain. These overlays indicate that there is none of the amine reactive dye in the FMO and the single colour indicates that the reactive dye is not shifting the fully stained sample.

*FMO controls*

Fluorescence minus one (FMO) control was used to set gates and determine amount of spectral spread between channels. Relevant examples are shown in Online Figures 6 and 7.

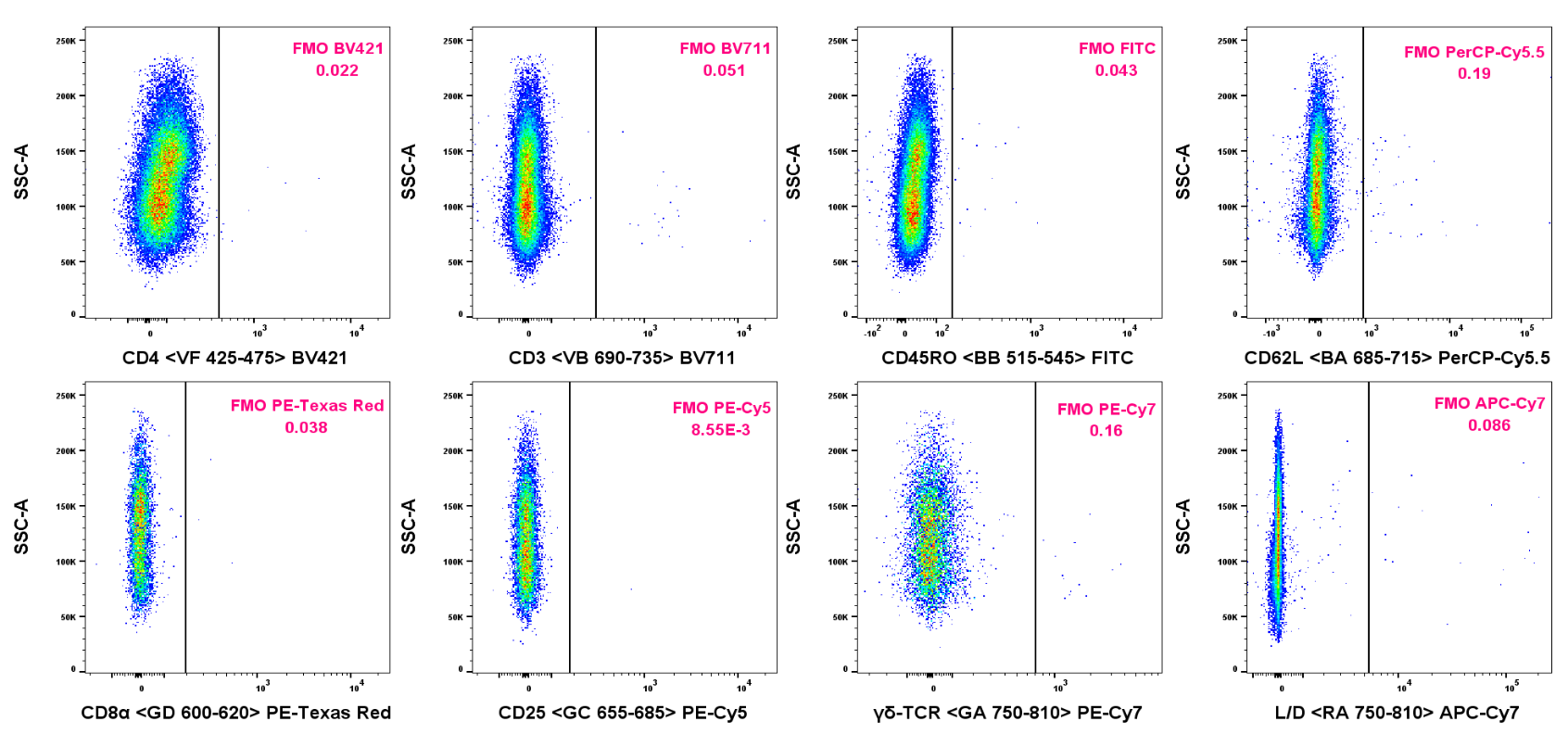

**Online Figure 7** Appropriate fluorescence intensity gating, using the fluorescence minus one control for all fluorochromes in this panel against the side scatter.

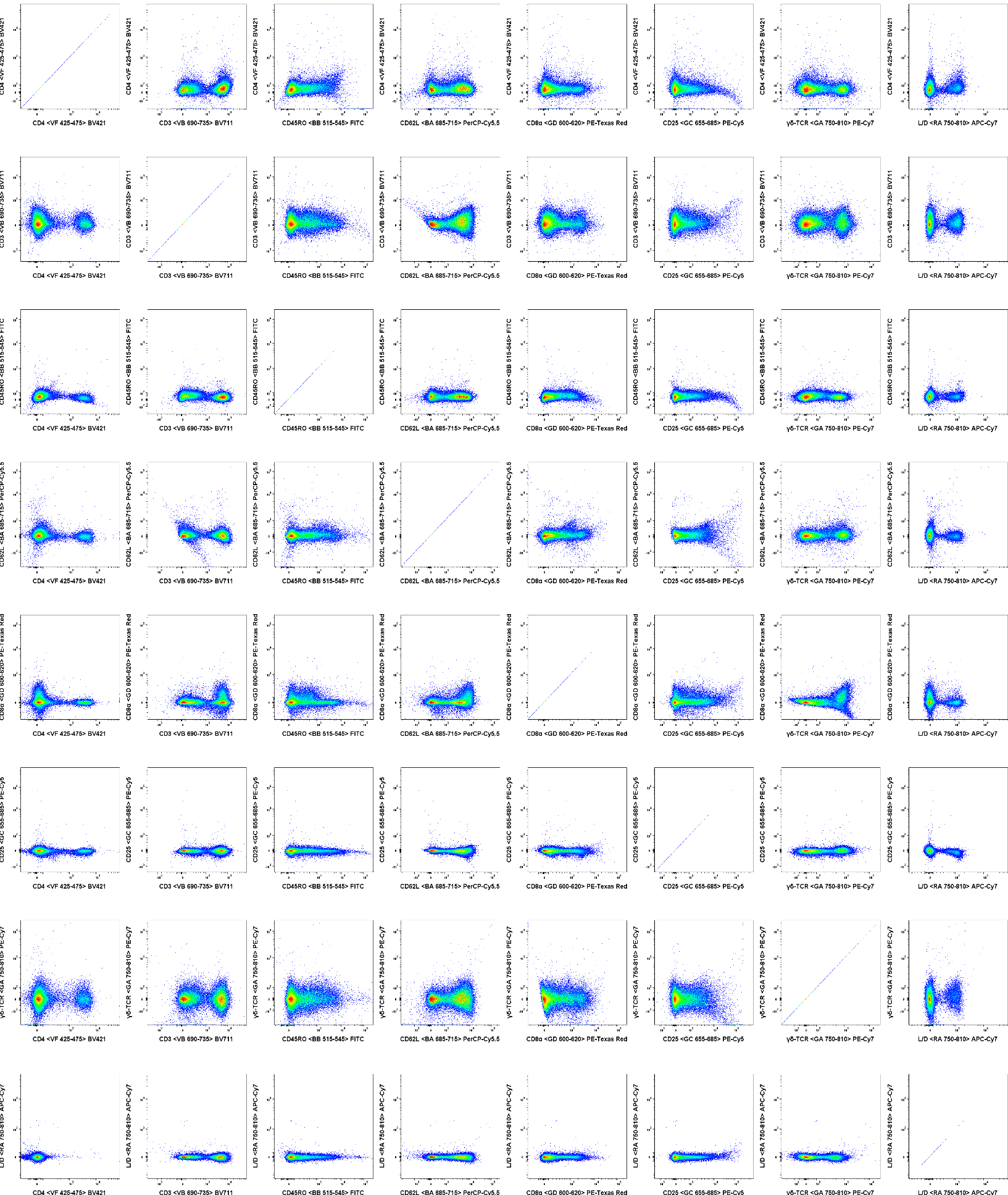

**Online Figure 8** Fluorescence minus one control matrix comparing all fluorochromes in this panel.

*Antibody titration*

To determine the optimal antibody concentration a 2-fold serial dilution was performed. This was done by adding 4ul of the 1mg/ml in-house conjugated antibody to 250ul of staining buffer (1:62.5 1:125, 1:250, 1:500, 1:1000, 1:2000, 1:4000, 1:8000 and an unstained sample) to determine the optimal concentration for each in-house conjugated antibody. The optimal concentration was picked as the lowest concentration with the highest resolution of the population of interest and having no or the least effect on the negative population (Online Figure 8). It is important to note that each new conjugate with the in-house conjugation kits needs to be titrated before being used in this panel and thus these dilutions are just representative of what was used in the final version of the panel.

**Online Figure 9** Two-fold dilutions were performed for each of the directly conjugated antibodies. The primary and biotinylated antibodies were titrated as above and both the secondary and the streptavidin-conjugated secondary were used at 1:3000. Titrated samples were concatenated and included an unstained sample to compare the negative population with the stained sample, due to background (indicated by the black line). Optimal dilutions for each antibody are indicated by the red box. Each new in-house conjugation was titrated before use during the development.

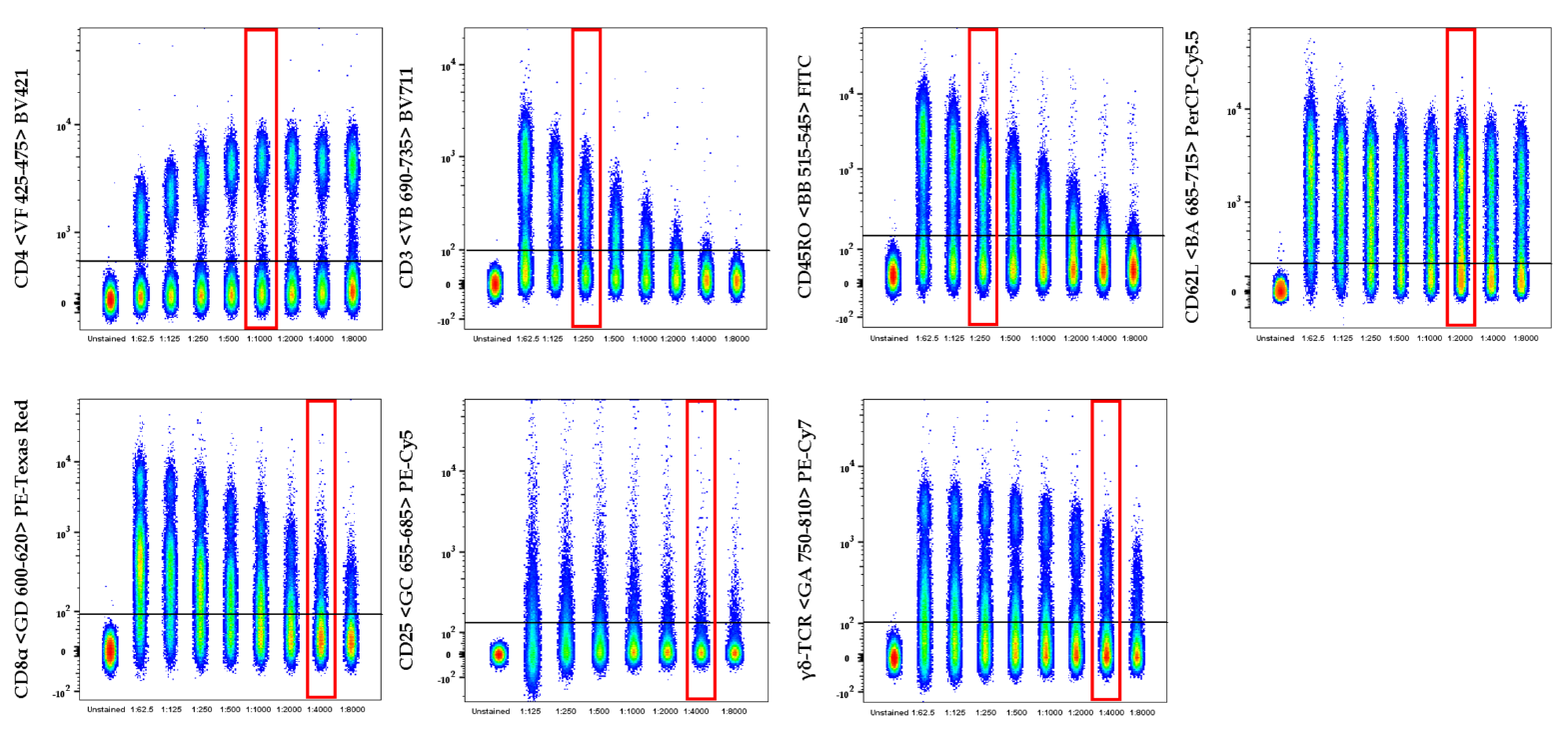

*Gating strategy*

The T cell subsets were discriminated using CD3 versus γδ-TCR, this allowed for the identification of unconventional T cells (Fig. 1). The unconventional T cells were further dissected with CD45RO and CD62L (Fig. 1). In parallel the CD3^+^γδ-TCR^-^ subset was interrogated by CD4 and CD8α to identify T helper and cytotoxic T cells, respectively. (Fig. 1). Thereafter the CD4 compartment was further characterised using CD25^+^ to identify activated T cells, whereas the CD25^-^ subset was further dissected using CD45RO and CD62L to determine the memory phenotype of the helper T cells (Fig. 1). The CD8α compartment was further characterised using CD45RO and CD62L to determine the memory phenotype of the cytotoxic T cells (Fig. 1). This gating strategy identifies 13 different T cell sub-populations (Fig. 1).

**Staining Protocol**

**Commercial materials:**

Corning™ Falcon™ 15mL Conical Centrifuge Tubes (VWR International, Cat No. 734-0452)

Horse serum 500 ml (Sigma, H1138)

PBSa (In-House supply)

Polystyrene 96 well conical ´V´ bottom plate (Elkay Laboratory Products (UK) Ltd., Cat No. MICR-TPV)

Staining buffer: PBSa with 0.5% Horse serum, 2mM EDTA (UltraPure™ 0.5M EDTA pH 8.0, Invitrogen™, Cat No. 15575020), 0.09% NaN_3_ (Sodium azide ReagentPlus^®^, Cat No. S2002-25G)

Autotube 1.1ml tubes (Elkay Laboratory Products (UK) Ltd., Cat No. 000-MICR-120)

**Cell seeding procedure:**

1. Add cells to 10ml of PBS containing 10% horse serum and PBS in 15ml Falcon tube for 3 min and spin at 300×*g* for 15 min at 4°C.
2. Re-suspended cells in PBS and aliquot 100μl to each well (96 V-well plate) so that each well contains 1×10^6^ cell/well.
3. Centrifuge plate at 300×*g* for 5 min at 4°C and supernatant removed (for all centrifugation steps below).

**Staining:**

4. Add 100μl of CD3 primary antibody at 1:1000 in cold staining buffer to the cells and resuspend by gently pipetting.
5. Incubate 15 min at RT in the dark.
6. Centrifuge and wash ×2 with 200μl of cold staining buffer.
7. Add 100μl of BV711 secondary antibody at 1:3000 in cold staining buffer to cells and, resuspend by gently pipetting.
8. Incubate 15 min at RT in the dark.
9. Centrifuge and wash ×2 with 200μl of cold staining buffer.
10. Add 100μl of conjugated antibody mix, at appropriate dilutions in cold staining buffer, to each sample well and resuspend by gently pipetting.
11. Incubate 15 min at RT in the dark.
12. Centrifuge and wash ×2 with 200μl of cold staining buffer.
13. Add 100μl of Streptavidin-BV421 antibody at 1:3000 in cold staining buffer to cells and, resuspend by gently pipetting.
14. Incubate 15 min at RT in the dark.
15. Centrifuge and wash ×1 with 200μl of cold staining buffer.
16. Centrifuge and wash ×2 with cold PBS only.
17. Add 100μl of LIVE/DEAD NIR diluted to 1:2000 in cold PBS to cells, resuspend by gently pipetting.
18. Incubate 10min at RT in dark.
19. Centrifuge and wash ×2 with cold PBS only.
20. Re-suspend pellet by gently pipetting 200μl cold PBS and transfer to 1.1ml flow tubes for analysis.
